# Reduced Auditory Perception and Brain Response with Quiet TMS Coil

**DOI:** 10.1101/2024.06.24.600400

**Authors:** David L. K. Murphy, Lari M. Koponen, Eleanor Wood, Yiru Li, Noreen Bukhari-Parlakturk, Stefan M. Goetz, Angel V. Peterchev

**Author notes:** Corresponding author Angel V. Peterchev, 40 Duke Medicine Circle Box 3620 DUMC, Durham, NC 27710 USA.

## Abstract

**BACKGROUND:** Electromagnetic forces in transcranial magnetic stimulation (TMS) coils generate a loud clicking sound that produces confounding auditory activation and is potentially hazardous to hearing. To reduce this noise while maintaining stimulation efficiency similar to conventional TMS coils, we previously developed a quiet TMS double containment coil (qTMS-DCC).

**OBJECTIVE:** To compare the stimulation strength, perceived loudness, and EEG response between qTMS-DCC and a commercial TMS coil.

**METHODS:** Nine healthy volunteers participated in a within-subject study design. The resting motor thresholds (RMTs) for qTMS-DCC and MagVenture Cool-B65 were measured. Psychoacoustic titration matched the Cool-B65 loudness to qTMS-DCC pulsed at 80, 100, and 120% RMT. Event-related potentials (ERPs) were recorded for both coils. The psychoacoustic titration and ERPs were acquired with the coils both on and 6 cm off the scalp, the latter isolating the effects of airborne auditory stimulation from body sound and electromagnetic stimulation. The ERP comparisons focused on a centro-frontal region that encompassed peak responses in the global signal.

**RESULTS:** RMT did not differ significantly between the coils, with or without the EEG cap on the head. qTMS-DCC was perceived to be substantially quieter than Cool-B65. For example, qTMS-DCC at 100% coil-specific RMT sounded like Cool-B65 at 34% RMT. The general ERP waveform and topography were similar between the two coils, as were early-latency components, indicating comparable electromagnetic brain stimulation in the on-scalp condition. qTMS-DCC had a significantly smaller P180 component in both on-scalp and off-scalp conditions, supporting reduced auditory activation.

**CONCLUSIONS:** The stimulation efficiency of qTMS-DCC matched Cool-B65, while having substantially lower perceived loudness and auditory-evoked potentials.

**Highlights:**

- qTMS coil is subjectively and objectively quieter than conventional Cool-B65 coil
- qTMS coil at 100% motor threshold was as loud as Cool-B65 at 34% motor threshold
- Attenuated coil noise reduced auditory N100 and P180 evoked response components
- qTMS coil enables reduction of auditory activation without masking

## 1. Introduction

Transcranial magnetic stimulation (TMS) uses electromagnetic induction to non-invasively stimulate, probe, and modulate activity in targeted cortical areas and wider brain networks in both clinical and basic research settings [1]. However, each TMS pulse produces mechanical vibration in the coil placed over the subject’s head that results in a loud clicking sound [2,3]. As it can reach peak sound pressure level (SPL) up to 140 dB(Z), exposure to the coil sound poses a risk to the hearing of TMS subjects and operators [4], and must be taken into account in safe clinical and research practices [5]. Even with proper hearing protection in place, TMS tolerability could be reduced in patient populations sensitive to sound [6–8]. The loud click also engages widely distributed auditory brain areas, as observed with positron emission tomography [9], functional magnetic resonance imaging [10,11], and electroencephalography (EEG) [12].

In TMS-EEG experiments, the concurrent auditory effects of TMS are particularly difficult to dissociate from its somatosensory and electromagnetic effects. Considerable effort has gone toward the segregation of a “true” TMS evoked potential (TEP) from the “unwelcome guest” [13] of auditory evoked potentials (AEP), somatosensory evoked potentials (SEP), and artifacts from scalp muscle activation [14]. The somatosensory components result from both the coil vibration on the scalp and the electromagnetic stimulation of scalp, facial, and ocular muscles close to the coil [15]. Some mitigation of the scalp pain due to coil vibration can be achieved by inserting a thin foam sheet between the coil and the subject’s head [16,17]. Approaches to suppress the AEP have included signal processing [18–22], auditory masking [23,24], and the use of various sham conditions that mimic the auditory and somatosensory stimulation arising from TMS [16,25,26].

These approaches to address the effects of TMS sound have yielded varying results [22,27,28]. The use of signal decomposition techniques in analyses may not preserve the temporal and spatial elements of the TEP [29]. Additionally, no single sham method independently removes auditory or somatosensory stimulation responses without altering the conditions of standard TMS use [16]: coil position, coil contact with the scalp, source of scalp stimulation, or source of auditory stimulation (air or bone conducted). Masking sound may influence the amplitude of motor evoked potentials (MEPs) elicited by TMS to primary motor cortex (M1), which provide a standard measure of cortical excitability [30]. Generally, nonlinear interactions between the sound and electromagnetically induced stimulation present a challenge for methods that aim to subtract computationally the effects of the sound.

To help mitigate the TMS coil vibration and sound, we previously developed a quiet TMS double containment coil (qTMS-DCC) [31]. qTMS-DCC suppresses the vibration of the coil winding with a dense inner casing and isolates it from the coil surface via an air gap with minimal structural connection between the winding case and the coil enclosure. qTMS-DCC was also designed to replicate the stimulation efficiency (induced electric field relative to device output) and inductance (determining the pulse duration) of the MagVenture Cool-B65 coil (MagVenture, Denmark) [31]. Acoustic measurements demonstrated that qTMS-DCC reduces the SPL of TMS pulses by 25 dB(Z) for matched estimated electromagnetic stimulation effect [31]. The practical utility of reduced acoustic output from TMS includes protecting the hearing of subjects and device operators, increasing the tolerability of treatment, reducing auditory stimulation without using sound masking, and enabling psychotherapy during TMS [32].

In this within-subject study in healthy human volunteers, we quantified the performance of qTMS-DCC and compared it to the standard MagVenture Cool-B65 coil. We verified that qTMS-DCC matches the stimulation efficiency of Cool-B65 with resting motor threshold comparisons and that qTMS-DCC is perceived as quieter during psychoacoustic testing. We then compared event-related potentials (ERPs) produced by each coil during active stimulation of M1 and auditory-only stimulation, considering differences in both the global signal and a centro-frontal region of interest (ROI). The experimental results corroborate the expected reduction in sound perception and acoustic brain activation with qTMS-DCC.

## 2. Methods

This study was approved by the Duke University Health System Institutional Review Board. Further details on methods are available in Supplementary Material.

### 2.1 Participants

Healthy participants (n = 9, 5 female) were recruited from the local population. The inclusion age range was 18–35 years to avoid confounds of age-related hearing loss, and the actual range was 19–29 years (23.5 ± 2.8 years, mean ± standard deviation). The participants were screened for normal hearing, use of psychoactive substances, and previous or ongoing neuropsychiatric disorders.

### 2.2 Experimental Conditions and TMS setup

Prior to the TMS experimental session, participants were scanned with a 3 T MRI machine to acquire T1-weighted anatomical images for neuronavigation. The TMS experimental session comprised two experiments separated by a short break (Table 1). The experiments compared the coil stimulation strength, subjective loudness, and ERPs of qTMS-DCC to MagVenture Cool-B65.

**Table 1.**
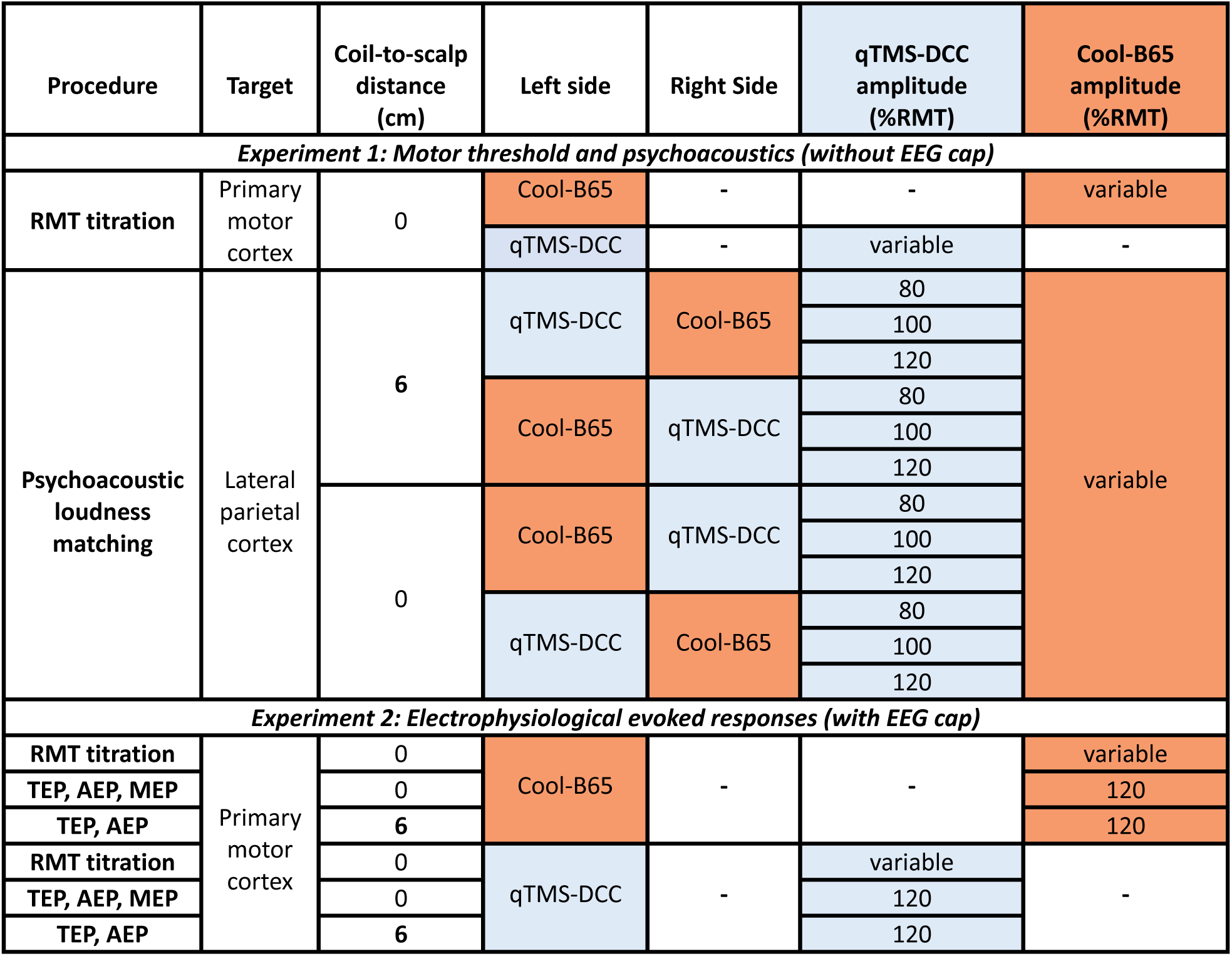
Summary of study design and experimental measurements.

During the experiments, the participants wore foam earplugs and were seated in a comfortable chair that also served as the base for the TMS coil holders and chin rest (Rogue Research, Canada). The complete experimental session, including preparation and testing, took approximately three hours per subject. Before and after, a brief TMS side effect rating scale and visual analog mood ratings were administered to evaluate side effects. qTMS-DCC and Cool-B65 were each connected to a separate MagPro X100 TMS device (MagVenture, Denmark) which delivered standard mode, biphasic pulses (300 µs period), with normal current direction during Experiment 1. During Experiment 1, RMT for each coil and subject was determined with the same MagPro X100. A single MagPro X100 was used for both coils in the EEG experiment (Experiment 2) to both determine RMT and deliver pulses during EEG recording. During all experiments, acoustic baffles (S2466x2 SORBER, ClearSonic) were placed around the MagPro devices to suppress the sound from the pulse generators and coil connectors. The qTMS-DCC and its cable (5.6 kg) were suspended by a custom counterweight system (see Supplementary Figure S1) to facilitate manipulation and to approximate the force placed on the scalp by a Cool-B65 coil (1.8 kg).

### 2.3 Stereotaxic neuronavigation

TMS coil position and orientation relative to the head were monitored with six degrees of freedom (three position and three orientation coordinates) using Brainsight (Rogue Research, Canada). For coil tracker calibration of qTMS-DCC, a custom calibration block was 3D-printed to mate the Brainsight calibration block to the center of the head-facing surface of qTMS-DCC. The anatomical T1-weighted MRI scan of each subject was imported into Brainsight, where scalp and cortical surface models were constructed for three-dimensional co-registration with the participant’s head in the experimental setting (see Supplementary Methods for full imaging details).

### 2.4 Electromyography

Electromyographic (EMG) recording of the right first dorsal interosseus (FDI) muscle was conducted with Ag/AgCl foam electrodes (Kendall 133, Covidien LLC, Ireland) on the skin and an EMG amplifier (BrainAmp ExG, Brain Products, Germany). Data from the BrainAmp ExG amplifier were passed to BrainVision Recorder software (BrainVision, USA) and then read by custom MATLAB (Mathworks, USA) programs that displayed the EMG data and performed online MEP analysis while recording the TMS pulse parameters. Trials that showed MEP activity of more than 50 μV peak-to-peak amplitude within the 100 ms interval immediately before the TMS pulse were marked as facilitated and excluded from the analysis.

### 2.5 Motor hotspot determination

The optimal stimulation location (hotspot) for each subject for inducing activity in the right FDI muscle was determined for both Cool-B65 and qTMS-DCC. The stimulator output started at 65% of the maximum stimulator output (MSO) with the coil placed tangential to the scalp over the hand knob area of the left motor cortex, with the coil oriented orthogonally to the motor strip (coil handle pointing backwards, angled ∼ 45° from midline). Single TMS pulses were then delivered with an interstimulus interval of ∼ 5 s, while moving the coil over a search grid. When registered to MNI space, the average Cartesian distance between the motor hotspot locations of the two coils were 2.16 ± 2.26 mm (range: 0.27–6.25 mm).

### 2.6 Resting motor threshold determination

Each subject’s resting motor threshold (RMT) for each coil was determined without and with an EEG cap on. RMT determination was guided by MTAT 2.0 software [33], which implements adaptive maximum-likelihood parametric estimation by sequential testing [34]. The initial stimulation intensity was set to the amplitude used in the motor hotspot search [35]. Because MTAT 2.0 can potentially misestimate RMT under certain conditions, ten additional pulses were delivered after MTAT 2.0 had converged on an initial RMT estimate.

### 2.7 Subjective loudness matching experiment

In Experiment 1, the coils were placed over the lateral parietal cortices on opposite sides. Participants then performed a two-alternative forced choice task designed to match the loudness of pulses delivered by Cool-B65 to pulses delivered by qTMS-DCC held at three fixed intensities (80, 100, and 120 %RMT).

Single pulses from each coil were delivered 400 ms apart, with the order of pulses from each coil pseudo randomly counterbalanced within sets of 10 sequential pulse pairs. The initial Cool-B65 pulse intensity was 80, 100, or 120 %RMT, matching that of qTMS-DCC. After the sequential stimuli were delivered, participants selected which pulse was louder (left or right) on a keyboard. Depending on the subject’s response, a custom MATLAB script raised or lowered the next stimulus intensity of Cool-B65. The intensity of the next Cool-B65 TMS pulse was calculated by a modified version of an algorithm designed to estimate equal loudness for two sound sources [36]. When participants changed their choice of which coil’s pulse was louder, this was considered a “reversal.” At the eleventh reversal, the titration was completed (see Supplementary Figure S2 for an example). On average, subjects performed eleven reversals after 20.0 ± 4.5 trials. The mean of the last six Cool-B65 reversal intensities was taken to be the level that best matched the loudness of qTMS-DCC. Participants performed two such runs for each qTMS-DCC intensity. The order of the fixed intensity presented in each run was randomized across the subjects.

This procedure was repeated for two separate coil-to-scalp distance conditions: with the coils on the subject’s head and with a coil-to-scalp distance of 6 cm. The latter condition effectively eliminated electromagnetic cortex and scalp stimulation as well as bone conduction of sound. The two coil-to-scalp distance conditions were then repeated after swapping the sides of the head on which the coils were placed, resulting in a total of twenty-four runs of the loudness matching algorithm. The initial scalp distance and the side on which the coils were placed were counterbalanced across participants.

### 2.8 Analyses of RMT and loudness matching

Differences in RMT between coils and EEG cap conditions were analyzed with a two-way repeated measures analysis of variance (RM-ANOVA). Analysis of the loudness matching paradigm utilized a three-way RM-ANOVA to identify differences between coil type, distance to scalp, and TMS intensity. The qTMS-DCC TMS pulse amplitudes and loudness-matched Cool-B65 TMS pulse amplitudes were divided by their respective mean RMT within condition to allow direct comparison of the amplitude adjustment for loudness matching between the coils. For the statistical analysis, the normalized TMS pulse amplitudes were log-transformed to reduce the heteroscedasticity of the distributions across the coils. To assess the robustness of the findings, we implemented leave-p-out cross-validation by repeating the RM-ANOVA on subsets of subjects, with all possible permutations of subsets ranging from three to eight subjects. The maximum p-value across all the permutations for each subset size was used to determined significance for the respective subset size.

### 2.9 EEG recording

In the second experiment, EEG signals after each stimulation were recorded to quantify the neural responses to qTMS-DCC versus Cool-B65. For this experiment, participants were fitted with an actiCap 64-channel EEG cap (layout M43-V1, Easycap GmbH, Germany) connected to an actiCHamp amplifier system (Brain Products, Germany). The cap was referenced to the Fz electrode and grounded at the Fpz electrode. EEG data were recorded with BrainVision Recorder (BrainVision, USA).

Foam padding (0.5 cm thick) was placed between electrodes on the EEG cap to reduce transmission of coil vibration to the EEG electrodes. The foam padding was also used during RMT titration. During EEG recording, the motor hotspot in left M1 was stimulated repeatedly at 120 %RMT.

All participants received 200–225 stimuli at 120 %RMT over the motor hotspot of the right FDI muscle; interstimulus interval was jittered between 1.5–2.5 seconds [37,38]. The same procedure was repeated with the coil 6 cm off the head to isolate AEPs. This resulted in two conditions for each coil in the TMS-EEG experiment, with the on-scalp condition producing active stimulation and off-scalp condition producing auditory stimulation alone. For each coil, both scalp distance conditions were performed in sequence to minimize the time between conditions and the disruption to neuronavigation and EEG cap setup. Counterbalance of the coil distance and coil type conditions (four conditions in total) was predetermined across participants.

### 2.10 ERP analysis

The following analyses were selected to compare responses to each coil, not to isolate and identify the complex interactions between somatosensory, auditory, and electromagnetic stimulation arising from TMS. All analyses were carried out in MATLAB using the toolboxes EEGLAB [39], Fieldtrip [40], and TESA [41]. Data between −8 ms and 15 ms relative to the TMS pulse were removed and interpolated before data were downsampled to 1 kHz (significance of results did not vary when performed on a subset of data sampled at 5 kHz).

Because we wished to compare the responses to each coil without specifically identifying components of sensory and TMS evoked potentials, we limited preprocessing to the following: automated channel removal via the EEGLAB function *pop_clean_rawdata,* epoch creation around TMS pulses (−1.25 s before to 1.25 s after the TMS pulse), bad trial removal with *pop_jointprob*, noise suppression via the SOUND algorithm [20] (*pop_tesa_sound*) during which electrodes were referenced to the global average, and bandstop (57–63 Hz stopband) and bandpass (1–100 Hz passband) filtering. Lastly, missing electrodes were spherically interpolated. After trial and channel rejection, 176–210 trials (192 ± 8) out of 200 to 225 trials, and 54–62 (58 ± 2) channels out of 63 were included. Trial epochs were trimmed to −400 ms before and 400 ms after TMS pulses, with average baseline signal for each electrode between −400 ms and −10 ms subtracted.

Peaks in the global mean field amplitude (GMFA) and region of interest (ROI) signals were extracted for all conditions, with nominal peak latencies and search windows corresponding to commonly reported TEP, AEP, and SEP components [16,25]. These components included the P30, N45, P60, N100, P180, and N280, all of which would appear as maxima in the GMFA signal. For each nominal peak, the mean GMFA latencies in all conditions were averaged across participants. These average peak latencies were then used to generate timepoints for the initial topographical analysis (Figure 3B–E).

After this, we selected a region of interest (ROI) from a group of electrodes with magnitude ≥ 2 µV in the grand average of all conditions at the mid- and late-latency peak timepoints from the GMFA signal (Figure 3). These timepoints were selected because response peaks were found for all subjects and conditions. This resulted in a centro-frontal ROI consisting of electrodes 1, 3, 6, 7, 35, 36, and 37, which are proximal to the Cz, FC1, FC3, FC2, F1, F2, and FC4 10-10 electrode locations, respectively. The significance of differences between conditions in GMFA, centro-frontal ROI signals, and topographical analyses was calculated with *fieldtrip*, using cluster-based permutation analyses to correct for multiple comparisons [40]. The time range for statistical analysis of the GMFA and ROI signals was 400 ms before the TMS pulse to 400 ms after. The mean signal between −400 and −10 ms relative to the TMS pulse was subtracted from each signal (baseline). Points between −10 ms and 20 ms relative to the TMS pulse were excluded from the analysis time range to negate variable effects of the TMS artifact removal between coils.

## 3 Results

3.1 Safety and side effects

The only reported side effects were mild headache and mild neck pain in one subject and mild neck pain in another subject. These side effects are common for TMS procedures [5]. None of the subjects reported hearing issues after TMS.

### 3.2 Stimulation efficiency

Within Experiments 1 and 2, the RMTs of the two coils were well matched (Figure 1 and Supplementary Table S1). In Experiment 1, without an EEG cap, the RMTs were 43 ± 10 %MSO for Cool-B65 and 42 ± 10 %MSO for qTMS-DCC (mean ± SD). In Experiment 2, with an EEG cap, the RMTs were 52 ± 10 %MSO for Cool-B65 and 53 ± 10 %MSO for qTMS-DCC. The RM-ANOVA disclosed no significant difference between the coils (F(1,8) = 0.740, p = 0.415). There were significant differences between the EEG cap conditions for each coil (F(1,8) = 320, p < 0.0001), as expected due to the coil-to-scalp distance added by the EEG cap. The interaction effect between the EEG cap condition and coil type was nonsignificant (F(1,8) = 5.26, p = 0.0509), indicating that the decay of the field away from the coils was comparable. The matched RMTs of the coils ensure that sound differences cannot be attributed to different energy levels delivered by the TMS devices.

**Figure 1.**
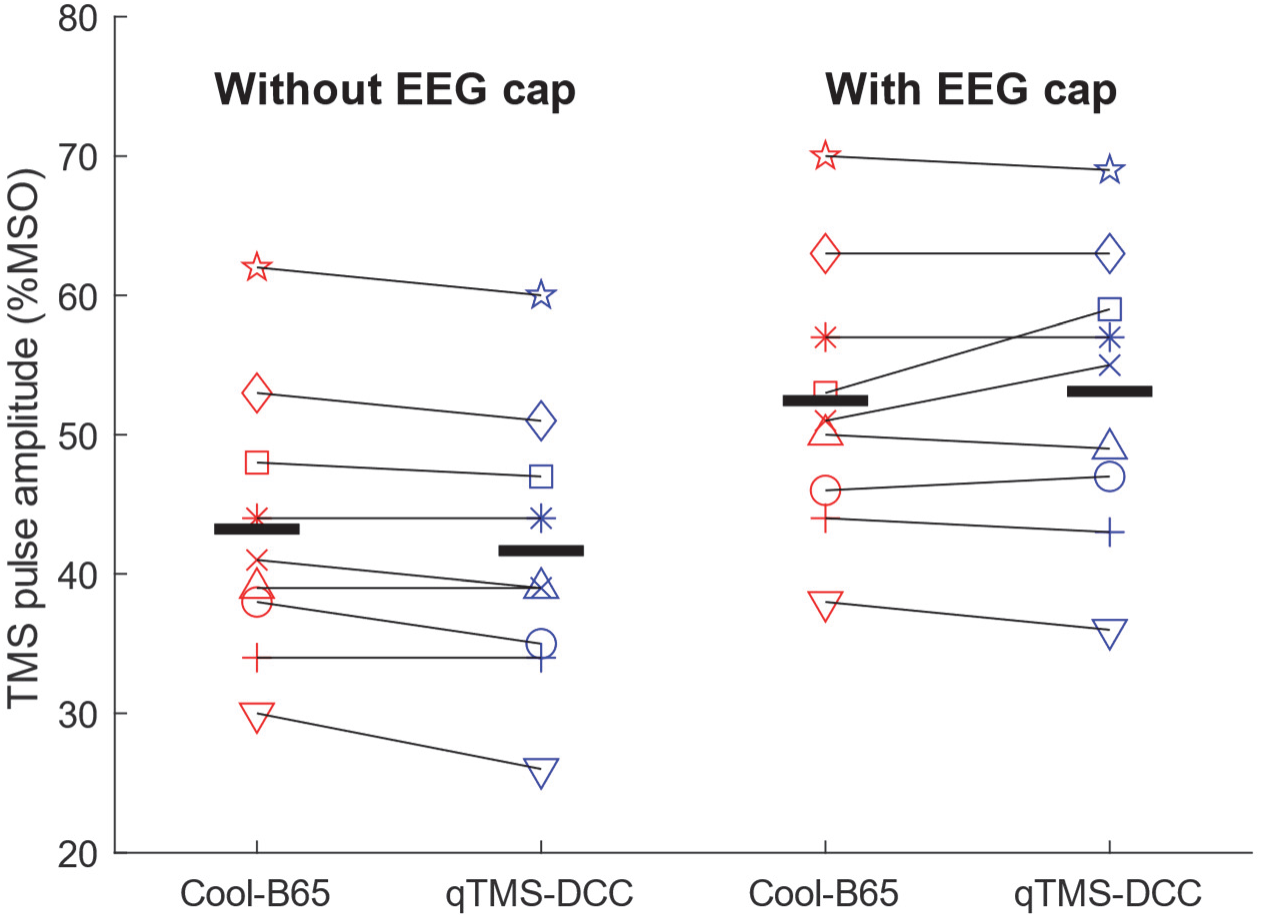
Resting motor threshold (RMT) as percentage of maximum stimulator output (MSO) for all subjects and each coil (Cool-B65 or qTMS-DCC) without (left) and with (right) EEG cap. Average RMTs displayed as thick horizontal lines. Between coil differences were not significant within the two EEG cap conditions (p = 0.415).

### 3.3 Perceived coil loudness

Figure 2 shows the TMS pulse amplitudes for the two coils corresponding to matched perceived loudness. In the coil-on-scalp condition (Figure 2A), participants lowered the TMS pulse amplitude of Cool-B65 to 29.7, 34.4, and 42.8 %RMT on average to match the loudness of qTMS-DCC held at 80, 100, and 120 %RMT pulse amplitude, respectively. For the coil-off-scalp condition (Figure 2B), these reductions were to 32.5, 44.3, and 55.8 %RMT, respectively. Across conditions, the TMS pulse amplitude for Cool-B65 was 60.2% lower relative to qTMS-DCC at matched loudness (F(1,8) = 651, p < 0.0001). As expected, the effect of the qTMS-DCC pulse amplitude setting relative to RMT, which served as the loudness reference, on the individual pulse amplitudes at matched loudness was also significant (F(2,8) = 329, p < 0.0001). The effect of coil distance was not significant (F(1,8) = 5.22, p = 0.0517), and neither were the interactions between the factors, indicating the differences between the coils were consistent across all conditions. Importantly, these results were very robust as the leave-p-out cross-validation indicated that significance was preserved for any combination of only 3 or more subjects in the analysis (Supplementary Table S2).

**Figure 2.**
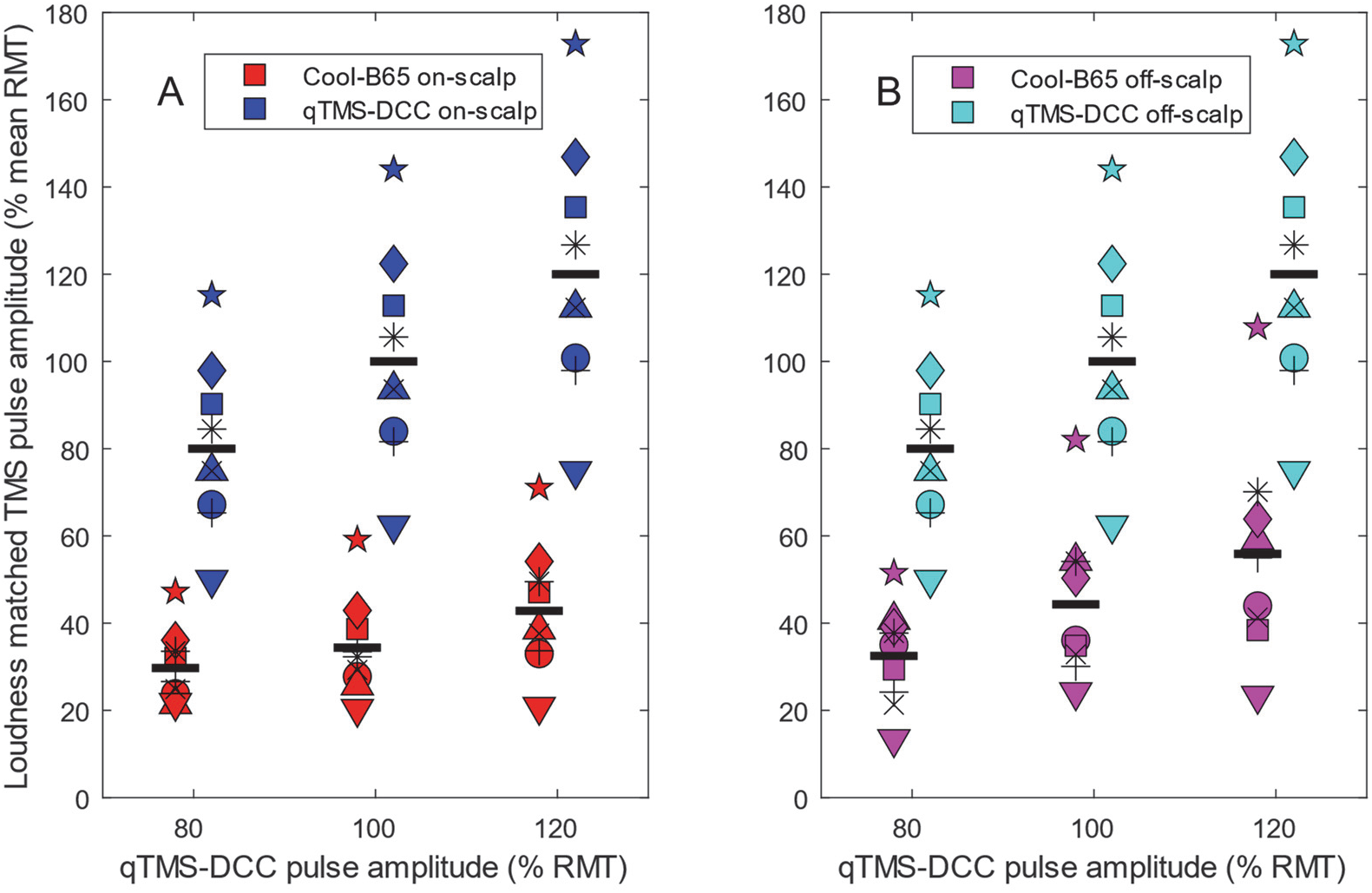
Perceived coil loudness matching. Pulse amplitude of qTMS-DCC was set to 80, 100, and 120 %RMT (x-axis), and subjects adjusted Cool-B65 pulse amplitude to match loudness. Resulting individual Cool-B65 and qTMS-DCC pulse amplitudes, normalized to mean RMT for each coil, are shown on y-axis for coils on scalp (A) and off scalp (B) with group averages denoted by black lines.

### 3.4 EEG evoked response potentials

#### Global mean field signal and response topography

In Figure 3 the GMFA responses and associated topographical maps show the general time course of evoked responses to all conditions. Averaged across all conditions, the latencies of GMFA maxima were 32, 56, 100, 186, and 275 ms (Figure 3A). Note that only peaks between 75–240 ms were detected in all subjects across conditions.

**Figure 3.**
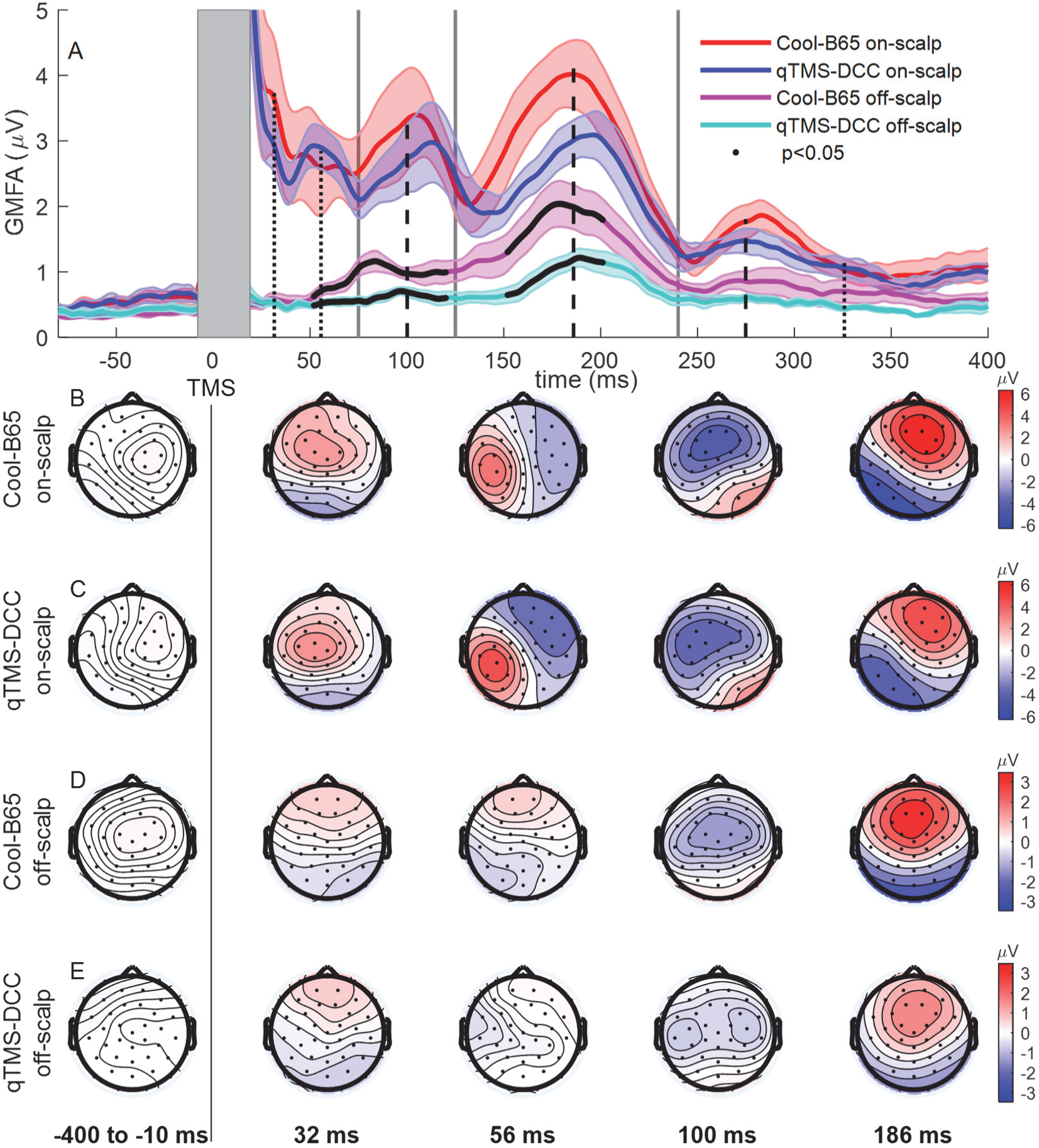
A. Global mean field amplitude (GMFA, grand-average ± SE) for coil on-scalp and off-scalp conditions. Average peak latencies across all conditions (black dashed lines for peaks found in all subjects, dotted lines for peaks found in some subjects). Signal from −9 ms to 19 ms (gray box) was excluded from consideration due to the TMS pulse artifact and artifact removal. Partitions between early-, mid-, and late-latency responses at 75, 130, and 240 ms are marked with vertical gray lines. Thick black traces indicate significant (p < 0.05) difference between coils within the off-scalp condition. B–E. Grand-average topographical plots at GMFA peak latency times.

In the on-scalp (active stimulation) condition, no significant difference between coils was detected in the GMFA signal, despite some apparent reduction in the GMFA signal of qTMS-DCC between 125 ms and 240 ms. For both coils, the responses at 32 ms to the active, on-scalp condition are seen at electrodes near the stimulation location, along the central sulcus, with the Cool-B65 producing slightly more frontal positivity (Figure 3B–C). The peak at 56 ms typically corresponds to somatosensory reafferent feedback. The response topography at 100 ms appears to be bilateral in centro-frontal electrodes for both coils. At 186 ms, the topography for both coils are very similar, with a centro-frontal and frontal response that is lateralized contralaterally to the stimulation side.

In the off-scalp (auditory-only) condition, Cool-B65 produced significantly larger GMFA signal around the 100 ms and 186 ms maxima. In the topographical plots (Figure 3D–E), Cool-B65 produces an observably stronger off-scalp response at 100 ms, with a more centralized distribution than that of the response to qTMS-DCC. The response to qTMS-DCC is symmetrically organized with peak negative responses closer to the auditory cortices in the temporal lobes. The most apparent similarities in response location appear at the 186 ms GMFA peak location, with qTMS-DCC producing a smaller response (as observed in the GMFA signal).

#### Centro-frontal ROI response

The electrodes selected by the magnitude of measured response all fell within the centro-frontal region (Figure 4). For these electrodes, the average ROI signal latencies across conditions had maxima at 31, 56, and 183 ms and had minima at 46 and 93 ms. The only minima and maxima located across conditions and for all subjects were in the mid- and late-latency ranges corresponding to the N100 and P180 components. Four post-stimulus time windows for topographic analyses of the stimulus responses (Figure 5B,D) were defined by the mean minimum and mean maximum peak latencies across all conditions for each nominal peak location.

**Figure 4.**
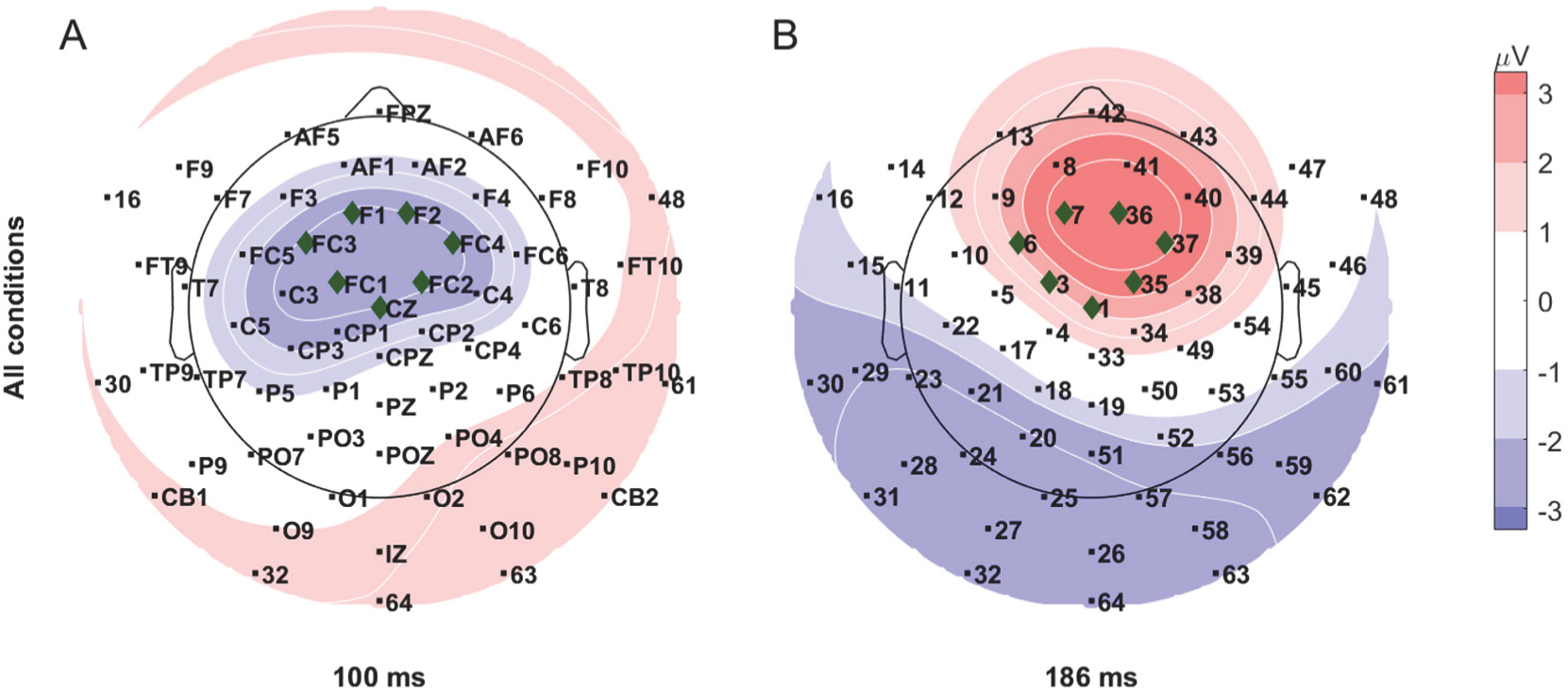
Centro-frontal region of interest (ROI) electrode selection and topography of grand-average response across all conditions (on-scalp and off-scalp for both coils). All ROI electrodes (green diamond markers) had absolute grand-average signal magnitude ≥ 2 μV at (A) mid-latency (100 ms) and (B) late-latency (186 ms) GMFA peaks which were detected for all subjects. Electrodes with letter labels are named after proximal 10–10 system electrode positions; remaining electrodes are numbered following the custom cap convention. Equipotential contour lines are shown in white.

**Figure 5.**
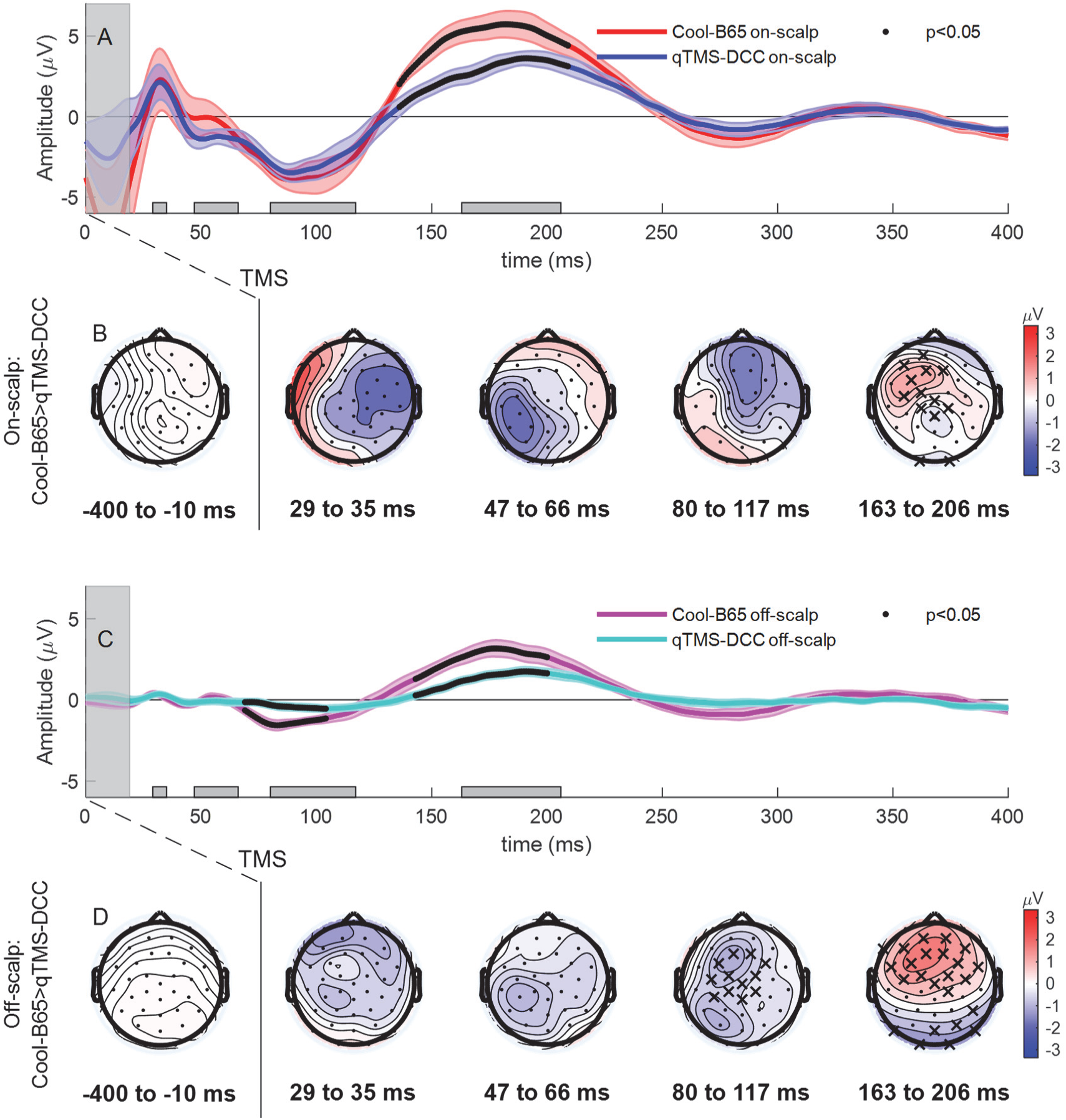
Grand-average signals from centro-frontal ROI electrodes (grand average ± SE) for on-scalp (A) and off-scalp (C) conditions. Thick black traces denote significant (p < 0.05) differences between the coils. Grand average topographic maps of response differences between coils from on-scalp (B) and off-scalp (D) conditions. Electrodes with significant (p < 0.05) differences between coils marked with “x” (negative differences in the blue areas and positive differences in the red areas). Individual traces for each subject are shown in Supplementary Figure S3.

In the on-scalp condition, the coils produced overlapping ROI signal waveforms until 125 ms (Figure 5A). qTMS-DCC had a significantly smaller P180 component than Cool-B65 in both on- and off-scalp conditions. The ROI signal differences between coils, in the on-scalp condition, are reflected in the scalp topography of the response between 163 ms and 206 ms (Figure 5B), where six out of the seven ROI electrodes show a significant decrease in response to qTMS-DCC.

During off-scalp stimulation, the centro-frontal ROI signal was not significantly different in the early-latency period. In the mid- and late-latency periods, significant differences in the N100 and P180 were observed. This was reflected in significant differences between all ROI electrodes in the 80–117 ms and the 163–206 ms time windows (Figure 5D).

## 4. Discussion

Compared experimentally to the conventional Cool-B65 coil, qTMS-DCC matched the stimulation efficiency, was rated by subjects as substantially quieter, and evoked less auditory brain activation.

### 4.1. Stimulation efficiency

The RMTs of the Cool-B65 and qTMS-DCC coils were well matched and increased similarly by approximately 10 %MSO with the EEG cap in place (Figure 1). This increase is expected from the addition of coil-to-cortex distance by the EEG cap and EEG electrodes, which was made consistent with foam padding between the EEG electrodes. The similar RMT and attenuation of induced electric field with coil-to-cortex distance were expected, as qTMS-DCC was designed to induce an electric field that matched that of Cool-B65 while minimizing coil heating and transmission of coil vibration [31]. The matched electrical performance of the coils helps to isolate auditory effect differences between the two.

### 4.2 Perceived loudness

The qTMS-DCC coil was perceived to be substantially quieter than the Cool-B65 coil. For example, active TMS with qTMS-DCC at coil-specific 100 %RMT sounded like Cool-B65 at 34 %RMT. These subjective ratings are consistent with our direct sound measurements showing an SPL reduction of about 25 dB(Z) between qTMS-DCC and Cool-B65 [31][4]. Notably, Cool-B65 is relatively quiet among commercial coils, and therefore the sound reduction for qTMS-DCC represents robust advantages compared to other commercial coils.

### 4.3 Brain response

Comparing Cool-B65 to qTMS-DCC with EEG recordings, the centro-frontal ROI ERP had larger magnitude of the P180 component for both on-scalp (active) and off-scalp (auditory only) TMS as well as the N100 component in the off-scalp TMS condition (Figure 5), indicating stronger auditory stimulation by Cool-B65. The auditory nature of this difference was further supported by the global brain response, as in the off-scalp condition the GMFA signal was significantly larger for Cool-B65 than qTMS-DCC from 50–200 ms (Figure 3A), which could only be attributed to air-conducted coil sound.

The first three TEP components have been frequently identified as the N15-P30-N45 complex: N15 at the stimulation site, more central P30, and parietal N45 [21,42]. While we use ERP component nomenclature that mirrors the timing of peaks in our data (P30, N45, P60, N100, P180), we recognize that the short latency components have been observed at a range of timepoints and have been referred to by different names (P25, N40, P55). The later P60 component has been observed in TMS-EEG studies of diverse populations and confirmed with optimized sham procedures [25,43]. During on-scalp stimulation of M1, both qTMS-DCC and Cool-B65 coil produced similar responses in the centro-frontal ROI, until the late-latency period (Figure 5A). The overlap of the P30 signal generated by each coil in the scalp topography maps (Figure 3B–C) and in the ROI signal (Figure 5A), marks the similarity in the immediate, cortical response to stimulation. Differences in the responses to each on-scalp coil can be seen at latencies corresponding to the N45 and P60 TEP components; these differences fall within the standard error of the signals produced by each coil and are therefore not significant in this sample. The N45 and P60 components are nearly coincidental to the P50 component evoked from auditory stimulation alone [21,44]. The interaction between the N45 and P60 of the TEP and the P50 of the AEP is a possible explanation for the small positive increase in the ROI signal from Cool-B65, although the N45 and P60 are more localized to the parietal cortex.

Compared to the earlier components of the ERP, the later N100 and P180 responses are more clearly associated with sensory and auditory responses [27,45]. The N100 response to active TMS mirrors what has been observed in previous studies [16,46]. The overlap between coils of the mid-latency ROI signals likely reflects how the N100 serves as a generic marker of somatosensory (afferent and reafferent) and auditory stimulation rather than as a marker of direct cortical excitation by TMS [47]. Importantly, during the off-scalp condition, the ROI and topographical analyses show significant between-coil differences for the ROI electrodes from N100 component (81–115 ms).

Considering the P180 component evoked by qTMS-DCC compared to Cool-B65, the results from the on-scalp stimulation parallel the reductions observed in the off-scalp condition. This supports the notion that the P180 component observed in TMS-EEG studies can be attributed to the auditory stimulus generated by each TMS pulse, while the N100 and earlier components are largely driven by somatosensory and electromagnetic stimulation. The significantly smaller P180 component from on-scalp qTMS-DCC stimulation demonstrates that late-latency AEP components are independently reduced by attenuation of the coil sound. That this can be achieved with qTMS-DCC without the addition of auditory masking helps unweave the overlapping AEP, SEP, and TEP signals associated with TMS-EEG measures.

These ERP results add a new perspective to the discussion of auditory stimulation effects from TMS, as we were able to reduce the late-latency P180 components of AEP during active stimulation, without the use of potentially confounding methods ranging from auditory masking to post-acquisition signal processing. Further investigation of the effects of reduced coil noise on early-latency responses to TMS [45] is merited but is beyond the scope of this study. Finally, we have demonstrated with SPL measurements that TMS sound can be suppressed further with the use of a novel ultra-brief-pulse generator, which could additionally reduce auditory activation [7,48,49], providing a direction for future investigation.

### Conclusions

The qTMS-DCC coil design substantially reduces perceived loudness and auditory activation of the brain while matching the stimulation efficiency of the commercial Cool-B65 coil. The reduction of auditory stimulation with qTMS-DCC could enhance safety, tolerability, blinding, and functional selectivity in clinical and research applications of TMS.

## Data availability

The data from this study are available in the National Institute of Mental Health Data Archive (NDA), https://nda.nih.gov, dataset identifier: 2531.

## Declaration of Interest

D.L.K.M., L.M.K, S.M.G, and A.V.P. are inventors on patents and patent applications on quiet TMS. L.M.K. has received patent royalties from Nexstim and consulting fees from Ampa Health. N.B.P serves on the medical and scientific advisory board of the Dystonia Medical Research Foundation. S.M.G. is inventor on patents on brain stimulation technology and has received research funding, royalties, or consulting fees from Magstim, Rogue Research, and Ampa Health. A.V.P. is an inventor on patents on TMS technology and has received equity options, scientific advisory board membership, and consulting fees from Ampa Health, patent royalties and consulting fees from Rogue Research, consulting fees from Magnetic Tides and Soterix Medical, equipment loan from MagVenture, and research funding from Motif. The other authors declare that they have no known competing financial interests or personal relationships that could have appeared to influence the work reported in this paper.

## Acknowledgements

Research reported in this publication was supported by the National Institute of Mental Health of the National Institutes of Health under Award Numbers R01MH111865 and R01MH127104. The study utilized TMS devices on loan from MagVenture Inc. The content is solely the responsibility of the authors and does not necessarily represent the official views of the funding agencies. The authors thank Dr. Sherri Smith for providing the audiometer and related training, R. Lane Daughtry for assisting with data sharing, and Dr. Risto Ilmoniemi for methodological advice.

## Notes

https://nda.nih.gov

