## Supplementary Materials for "Reduced Auditory Perception and Brain Response with Quiet TMS Coil"

#### Supplementary Material

David L. K. Murphy[1], Lari M. Koponen[1], Eleanor Wood[1], Yiru Li[1], Noreen Bukhari-Parlakturk[2],

Stefan M. Goetz[1,3,4,5], Angel V. Peterchev[1,3,4,6,*]

1. Department of Psychiatry and Behavioral Sciences, Duke University School of Medicine

2. Department of Neurology, Duke University School of Medicine

3. Department of Electrical and Computer Engineering, Duke University

4. Department of Neurosurgery, Duke University School of Medicine

5. Department of Engineering, Technical University Kaiserslautern

6. Department of Biomedical Engineering, Duke University

### S1 Supplementary Methods

#### S1.1 Subject screening
This screening includes the standard pure-tone threshold audiometry for normal hearing (Maico MI-25 screening audiometer), TASS form for eligibility to receive TMS, a survey for eligibility based on neurological function, DSM-IV MINI for eligibility based on psychiatric status, urine toxicology test for substance use or dependence, urine hCG test for pregnancy in women of child-bearing potential, and the Duke Brain Imaging and Analysis Center (BIAC) screening form for MRI scan eligibility. All subjects passed the audiometry tests for normal hearing before the TMS sessions.

#### S1.2 MRI acquisition
Prior to the TMS procedures, participants underwent an MRI imaging session (General Electric UltraHigh Performance MRI scanner, B0 field strength = 3 tesla, upgraded gradient coils) during which a structural, T1-weighted image was obtained (echo-planar sequence: voxel size = 0.5 $mm^3$, TR = 2304.66 ms, TE = 3.2 ms, flip angle = 8°, FOV = 256 $mm^2$, bandwidth = 31.3 Hz/pixel, 166 slices). While not used in this study, T2-weighted and diffusion-weighted scans were obtained as well (T2-weighted scan parameters: echo-planar sequence with fat saturation: voxel size = 0.8594 × 0.8594 × 1.5 $mm^3$, TR = 6.93 s, TE = 59.7 ms, flip angle = 90°, FOV = 220 $mm^2$, bandwidth = 281.3 Hz/pixel. Diffusion-weighted scan parameters: acquisition matrix = 144 $mm^2$, voxel size = 0.856 × 0.856 × 1.5 $mm^3$, TR = 46.6 s, TE = 64.7 ms, flip angle = 90°, FOV = 220 $mm^2$, bandwidth = 281.3 Hz/pixel, matrix size = 1282, B-value = 3000 s/$mm^2$, diffusion directions = 90).

#### S1.3 TMS equipment
The qTMS-DCC coil was supported by a commercial camera jib (ProJib) counterweighted with 40 lbs (Figure S1).

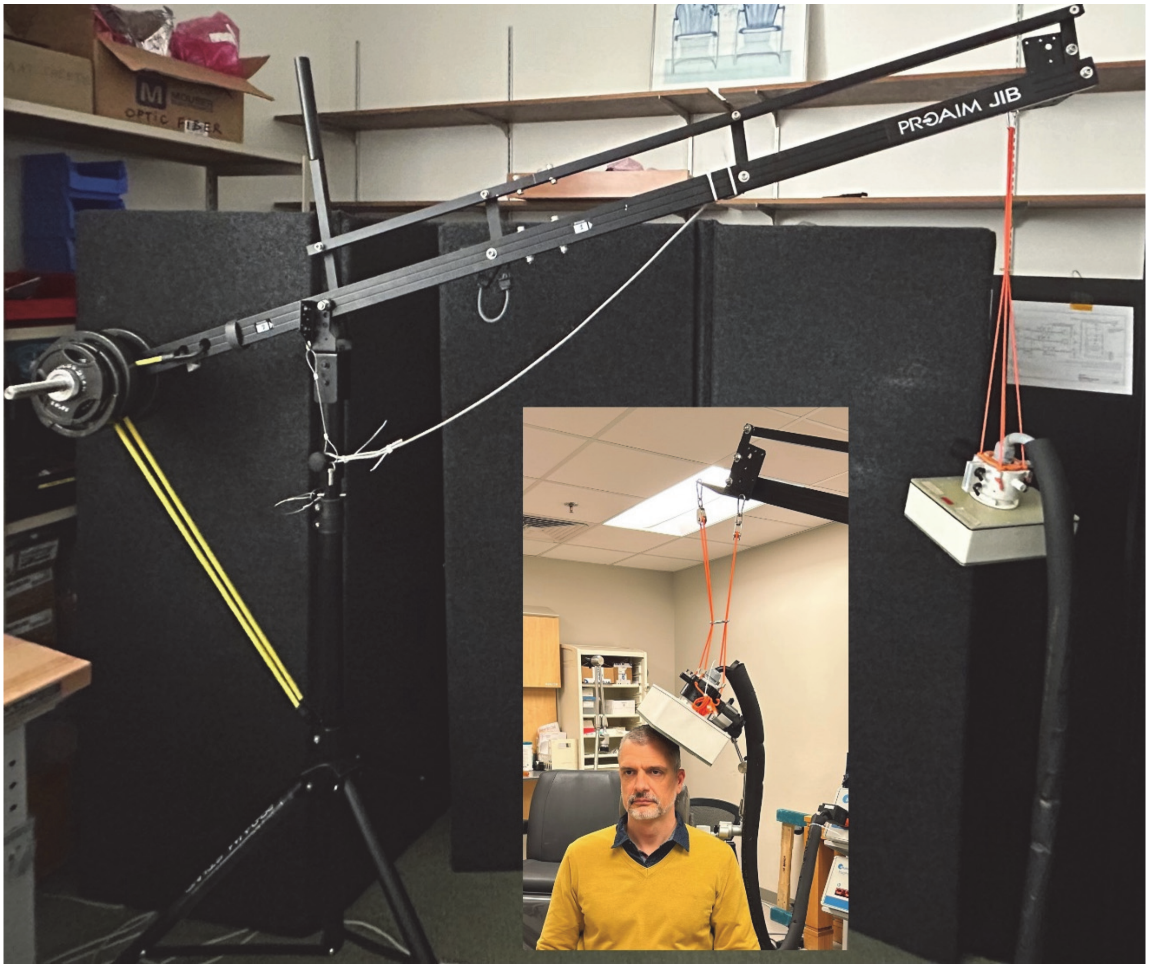


Figure S1. Counterweight system and example of the reduced weight of the coil on a subject's head.

#### S1.4 Electromyography and MEP analysis

***Skin Preparation:*** Prior to electrode placement, the electrode locations on the skin were cleaned with an isopropyl alcohol wipe, scrubbed with electrolytic paste, and wiped clean of excess paste.

***Signal acquisition:*** The BrainAmp ExG amplifier sampled with 16-bit resolution at 5000 Hz and bandpass filtered the EMG signal between 0.1 Hz and 1000 Hz and a 60 Hz bandstop mains filter. A trigger pulse from the MagPro X100 device was sent via BNC cable to BrainVision recorder via the amplifier which initiated the collection and display of EMG data from −200 ms before the trigger to 200 ms after the trigger.

***Signal processing:*** After online data transfer from BrainVision Recorder to our custom MATLAB app, the MEP amplitude was defined as the difference between the maximum and minimum values of the processed EMG waveform occurring between 20 ms and 60 ms after the TMS stimulus.

#### S1.5 Motor hotspot mapping

Single TMS pulses were then delivered with an interstimulus interval (ISI) of ~ 5 seconds. If MEP amplitude regularly exceeded 500 mV at the initial coil location, the stimulus intensity was lowered by 2% MSO. The coil was then moved in a search pattern of ~ 5 mm steps, moving the coil first in the inferior and superior (I–S) directions, then in the anterior and posterior (A–P) directions, and then in a radial pattern between the I–S and A–P axes. The location that produced the largest MEPs was marked in Brainsight and then coil orientation was optimized to produce the largest and most regular MEPs (1).

#### S1.6 Subjective loudness matching

For each qTMS-DCC pulse intensity, participants performed the loudness matching task twice. Cool-B65 stimulus intensity was controlled through a serial connection between the pulse generator and the task computer using a custom toolbox. An example of the titration steps of two runs are shown in Figure S1.

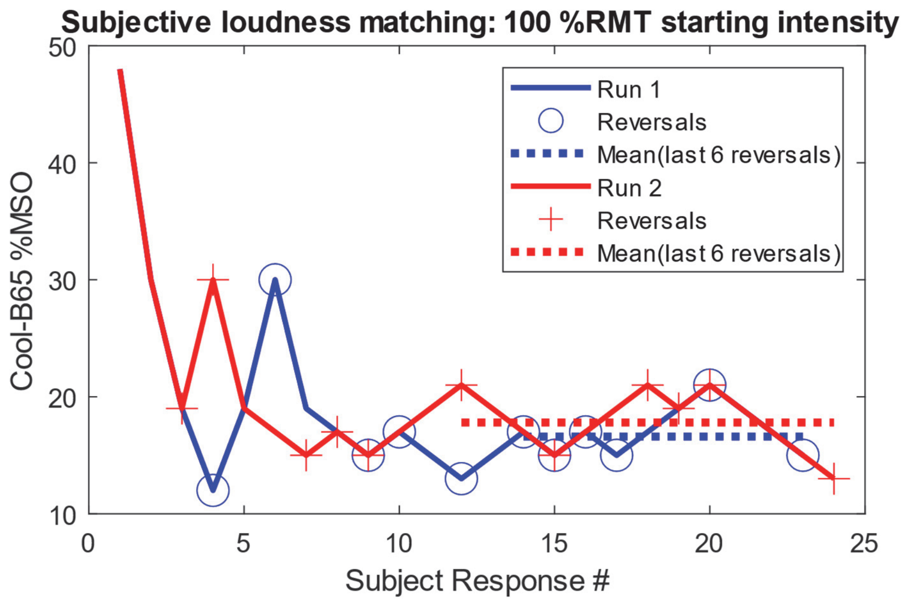


Figure S2. Example of subject adjusting Cool-B65 intensity to match qTMS-DCC loudness at 100 %RMT: 1st run (blue) and 2nd run (red) both starting at 100 %RMT of Cool-B65.

### S2 Supplementary results

Table S1. Individual resting motor thresholds for the two TMS coils (Cool-B65 and qTMS-DCC) without and with EEG cap.

| Subject | Cool-B65 | qTMS-DCC | Cool-B65 (EEG cap) | qTMS-DCC (EEG cap) |
|---|---|---|---|---|
| **S01** | 38 | 35 | 46 | 47 |
| **S02** | 34 | 34 | 44 | 43 |
| **S03** | 39 | 39 | 50 | 49 |
| **S04** | 48 | 47 | 53 | 59 |
| **S05** | 30 | 26 | 38 | 36 |
| **S06** | 62 | 60 | 70 | 69 |
| **S07** | 53 | 51 | 63 | 63 |
| **S08** | 44 | 44 | 57 | 57 |
| **S09** | 41 | 39 | 51 | 55 |
| **Mean** | 43.2 | 41.7 | 52.4 | 53.1 |
| **Std. dev.** | 9.9 | 10.1 | 9.8 | 10.3 |

Table S2. RM-ANOVA maximum p-values across all subject permutations for given subsample of subjects (leave-p-out cross-validation) for subjective loudness matching. Significant effects ($p < 0.05$) are marked in bold.

| | **Number of subjects** | | | | | | |
|---|---|---|---|---|---|---|---|
| **Factor** | **9 (all)** | **8** | **7** | **6** | **5** | **4** | **3** |
| **Coil** | **0.0000** | **0.0000** | **0.0000** | **0.0000** | **0.0001** | **0.0013** | **0.0111** |
| **Dist** | 0.0517 | 0.1062 | 0.2036 | 0.4023 | 0.8429 | 0.9016 | 0.9388 |
| **Intensity** | **0.0000** | **0.0000** | **0.0000** | **0.0000** | **0.0000** | **0.0001** | **0.0019** |
| **Coil × Dist** | 0.0517 | 0.1062 | 0.2036 | 0.4023 | 0.8429 | 0.9016 | 0.9388 |
| **Coil × Intensity** | 0.7600 | 0.9666 | 0.9991 | 0.9918 | 0.9998 | 0.9892 | 0.9922 |
| **Dist × Intensity** | 0.0906 | 0.2704 | 0.4911 | 0.7420 | 0.9142 | 0.9456 | 0.9314 |
| **Coil × Dist × Intensity** | 0.0906 | 0.2704 | 0.4911 | 0.7420 | 0.9142 | 0.9456 | 0.9314 |

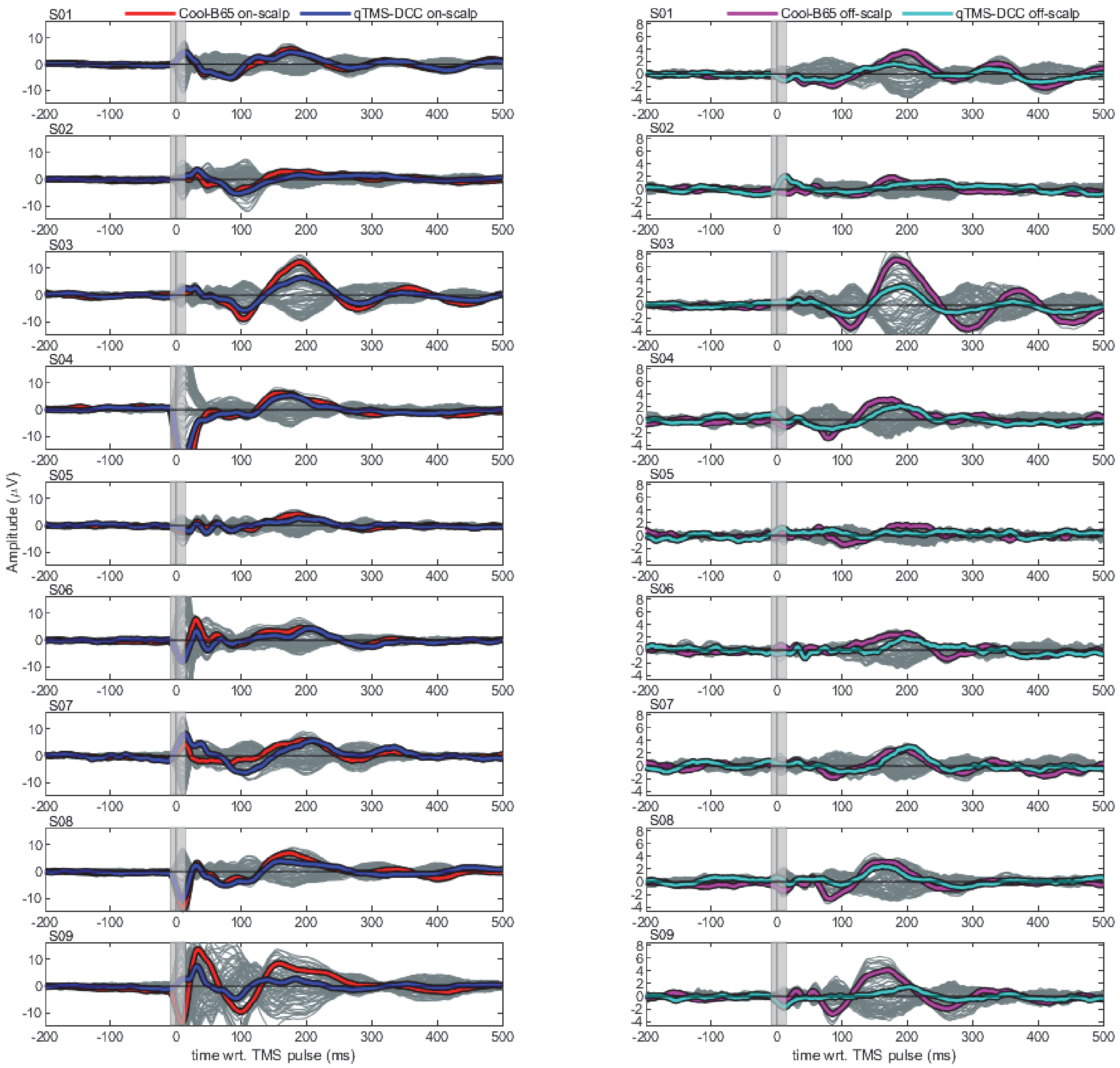


Figure S3. Individual butterfly plots for each subject and each condition in the centro-frontal ROI. Average signals of all trials for each of the 63 individual electrodes are shown as gray lines. Average ROI signal of all trials across electrodes is shown with colored lines. Period between −8 and 15 ms around TMS pulse is excluded from analysis due to TMS artifact and marked with a gray rectangle.
